# Genomic, transcriptomic and phenomic variation reveals the complex adaptation of modern maize breeding

**DOI:** 10.1101/005751

**Authors:** Haijun Liu, Xiaqing Wang, Marilyn L. Warburton, Weiwei Wen, Minliang Jin, Min Deng, Jie Liu, Hao Tong, Qingchun Pan, Xiaohong Yang, Jianbing Yan

## Abstract

The temperate-tropical division of early maize germplasm to different agricultural environments was arguably the greatest adaptation process associated with the success and near ubiquitous importance of global maize production. Deciphering this history is challenging, but new insight has been gained from the genomic, transcriptomic and phenotypic variation collected from 368 diverse temperate and tropical maize inbred lines in this study. This is the first attempt to systematically explore the mechanisms of the adaptation process. Our results indicated that divergence between tropical and temperate lines seem occur 3,400-6,700 years ago. A number of genomic selection signals and transcriptomic variants including differentially expressed individual genes and rewired co-expression networks of genes were identified. These candidate signals were found to be functionally related to stress response and most were associated with directionally selected traits, which may have been an advantage under widely varying environmental conditions faced by maize as it was migrated away from its domestication center. It’s also clear in our study that such stress adaptation could involve evolution of protein-coding sequences as well as transcriptome-level regulatory changes. This latter process may be a more flexible and dynamic way for maize to adapt to environmental changes over this dramatically short evolutionary time frame.

## Introduction

Maize (*Zea mays* ssp. mays) is essential to the global food supply, with current total maize grain production higher than any other crop (USDA FAS 2013). Maize is also used as a model to investigate crop evolution and improvement (Doebley et al. 2006). It is thought to have been domesticated from teosinte (*Zea mays* ssp. *parviglumis*) about 9,000-10,000 years ago in southwestern Mexico, which is a mid- to lowland tropical growing environment (Matsuoka et al. 2002; Van Heerwaarden et al. 2011). The remarkable conversion of a Mexican annual grass species into the top food, feed and industrial crop in the world resulted from the spread of temperate maize over several thousand years from its tropical geographic origin to the north and east across North America and to the south across most of Latin America, eventually creating a maize distribution from ∼40°S in Chile to ∼45°N in Canada (Matsuoka et al. 2002). Centuries ago, maize cultivation expanded further to East Asia, Europe, and Africa, and the temperate-tropical division remains in all crop-growing continents today. When faced with widely varying temperate conditions in temperature, day length, and disease susceptibility, maize adapted remarkably well. One major goal of adaptation studies is to identify specific genomic changes contributing to advantageous phenotypic performance in varying environmental conditions.

In order to identify the genetic factors driving maize evolution, researchers have explored a number of methods to reveal footprints of selection within the genome (Chia et al. 2012; Hufford et al. 2012; Jiao et al. 2012). It is intriguing that these changes occurred within such a short evolutionary time frame. The importance of transcriptional regulation of rapid phenotypic evolution has been a central tenet of recent studies (Swanson-Wagner et al. 2012; Carroll 2008; Ecker et al. 2012; Koenig et al. 2013). Genes with differential expression and altered expression networks could provide evidence of the contribution of transcriptome regulational changes to the adaptation process. RNA-seq (Wang et al. 2009) allows cost-effective exploration of both sequence and transcriptional variation, particularly in large and repetitive sequence-rich genomes such as maize.

Seed development, a critical process to both plant propagation and food supply, is a time in which DNA methylation and chromatin remodeling, and thus transcriptional patterns, are reshaped for the new generation (Ahmad et al. 2010; Wollmann et al. 2012; Zanten et al. 2011). Transcriptional variation may thus heavily influence seed-related traits via environmentally-sensitive epigenetic control (Zhang and Ogas 2009), which will be expressed as selectable variation throughout the lifetime of the plant (Kapazoglou et al. 2013; Casas et al. 2012). Most maize genes are expressed in seed or embryos, many of which are not expressed again (Cho et al. 2012; Sekhon et al. 2011). Thus, the seed offers the best window into visualizing differences that may account for adaptation.

To study the nature of maize adaptation from tropical to temperate growing regions, a panel of 368 diverse maize inbred lines (Li et al. 2013) (Supplemental Table S1) was characterized. We combined RNA-seq of seeds (15 days after pollination; Fu et al. 2013) with data from the MaizeSNP50 BeadChip, resulting in over one million high-quality SNPs and expression data from 28,769 genes, analyzed together with 662 phenotypic traits. These included morphological, agronomic, physiological and metabolic traits, many of which are also known to be important in stress adaptation (Bohnert and Sheveleva 1998; Bhargava and Sawant 2013). This study is the first systematic exploration of the mechanisms of the maize adaptation process with the goal of answering several specific questions: What phenotypic changes in temperate lines convey an advantage in novel environments? Which genomic regions were selected during the adaptation process? What phenotypes do these regions likely affect? To what extent do regulatory changes contribute to evolution? What beneficial value do they provide in the relatively short evolutionary time frame suggested in this study? Although only one organ (the seed) was sequenced, such knowledge will position us with general understanding of the maize adaptation process, and provide resources for developing breeding strategies to help corn producers cope with an increasingly erratic climate.

## Results

### Population level differences between temperate and tropical lines

The population-scaled recombination rates (ρ) in temperate and tropical lines were 1.078/kb and 2.644/kb, respectively. This is a reflection of different rates of LD decay, which was much faster in the tropical lines at the whole genome level (Supplemental Fig. S1). Recombination rate differences in temperate vs. tropical lines was smaller than the decrease in ρ seen in a previous study by Hufford et al. (2012) in landraces compared to teosinte (59% vs. 75%). A cross population composite likelihood approach (Chen et al. 2010) (XP-CLR) was used to identify extreme allele frequency differentiation over linked regions when comparing temperate to tropical subpopulations. We identified 701 regions containing 1660 selected genes at the highest 10% of XP-CLR values (Fig. 1, Supplemental Table S2,3), ranging in size from 10kb to 2,320kb, with an average of 150.9kb; this is shorter than the 322kb average region associated with domestication (Hufford et al. 2012). The combined length of selected regions was 105.7Mb, covering 5.2% of the genome. The selection coefficient in the adaptation process was 0.090, which is higher than domestication (0.015) and improvement (0.003) (Hufford et al. 2012), indicating a stronger selection pressure during the adaptation process. However, the coefficient and size of the genomic region associated with selection may not be directly comparable with adaptation, since most polymorphisms in the current study are based on expressed genes, compared to random sequence polymorphisms measured by Hufford et al. (2012)

**Figure 1.**
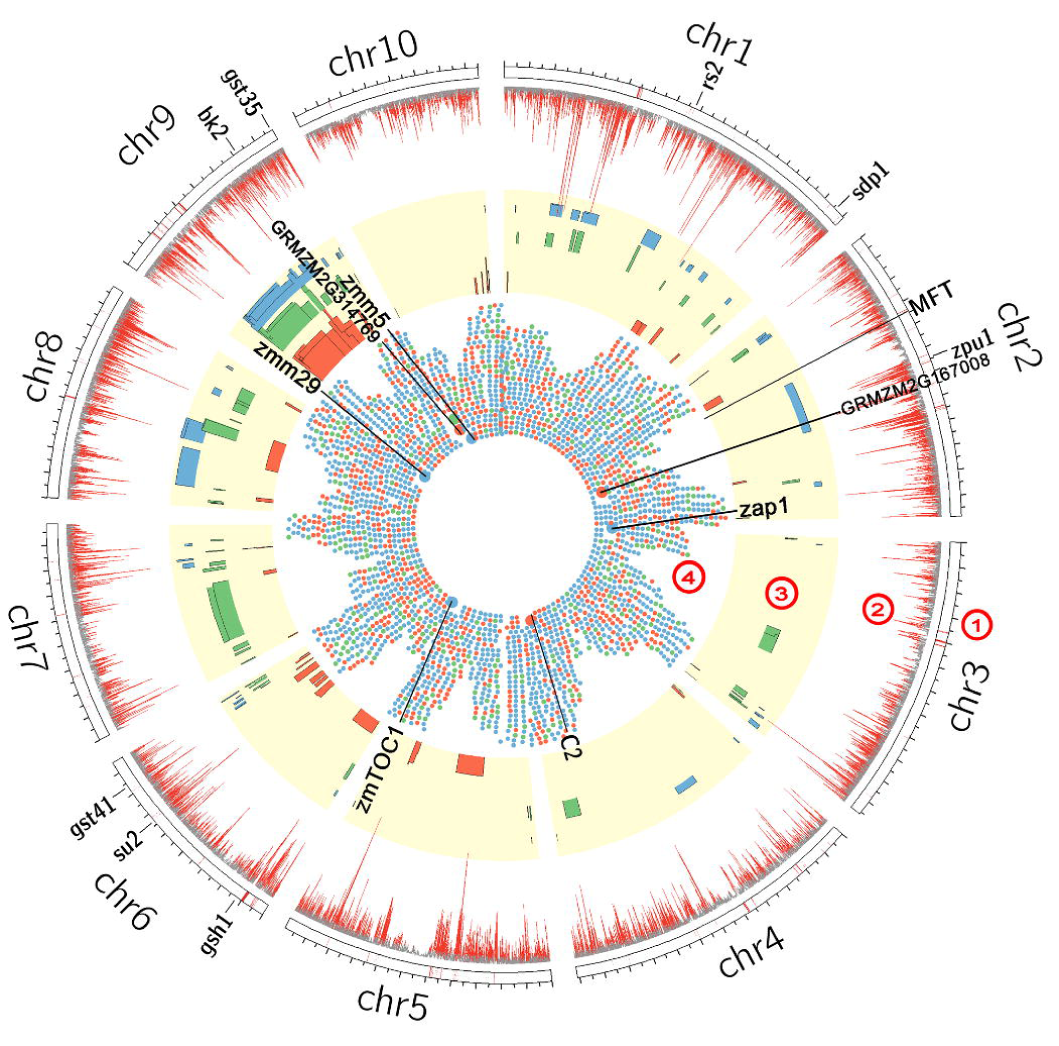
Integrated results of genome-selection (XP-CLR), transcriptome analysis (DE and AEC) and QTL mapping. ① Ten chromosomes of maize. ② XP-CLR value: the top 20% are marked in red, and the bottom 80% in grey. ③ QTL mapping results for days to silking (DTS, red), days to tasseling (DTT, green) and days to anthesis (DTA, blue). ④ Results of DE and AEC. Red and blue boxes indicate up- and down-regulated genes in temperate maize (TEM) relative to tropical (TST), respectively; green boxes indicate AEC genes. Some of the larger boxes are genes referred to in the text.

Nucleotide diversity (π) in the selected features identified by XP-CLR in temperate vs. tropical lines was 8.22E-04 and 8.42E-04, respectively, indicating a reduction of 2.4% in temperate lines. This decrease is less than the reduction of nucleotide diversity in selected features identified during domestication (17%) by Hufford *et al*. (2012). *F_st_* of selected features was 0.027, compared to 0.11 between teosinte and landraces, possibly due to the shorter time for adaptation from tropical to temperate than for domestication. However, this result is similar to 0.02 reported between landraces and improved lines (Hufford et al. 2012).

The divergence time between temperate and tropical subpopulations is of interest and can be associated with the development of agriculture and the spread of human civilization in the Americas; however, archeological information on this topic is incomplete and occasionally contradictory (Piperno and Pearsall 1998; Staller and Thompson 2002; Blake et al. 2006; Grobman et al. 2012). We proposed three models (detailed in the methods section, Fig. 2A–C) to estimate the time of divergence, resulting in the estimation of 3,400–6,700 years BP. This time frame is supported by recent archeological evidence (Haas et al. 2013) and implies that after domestication, maize cultivation rapidly expanded to temperate America (Fig. 2D). The molecular evidence thus suggests that improvement and adaptation here may not have been sequential and discrete processes, but overlapped in maize. Although gene flow between maize and its wild relatives has been shown to be of adaptive importance for maize evolution (Van Heerwaardena et al. 2011; Hufford et al. 2013), and has been measured in tropical maize (Warburton et al. 2011), it has been difficult to measure the rate or effect of gene flow in temperate lines following divergence; thus, we do not factor it into our analysis.

**Figure 2.**
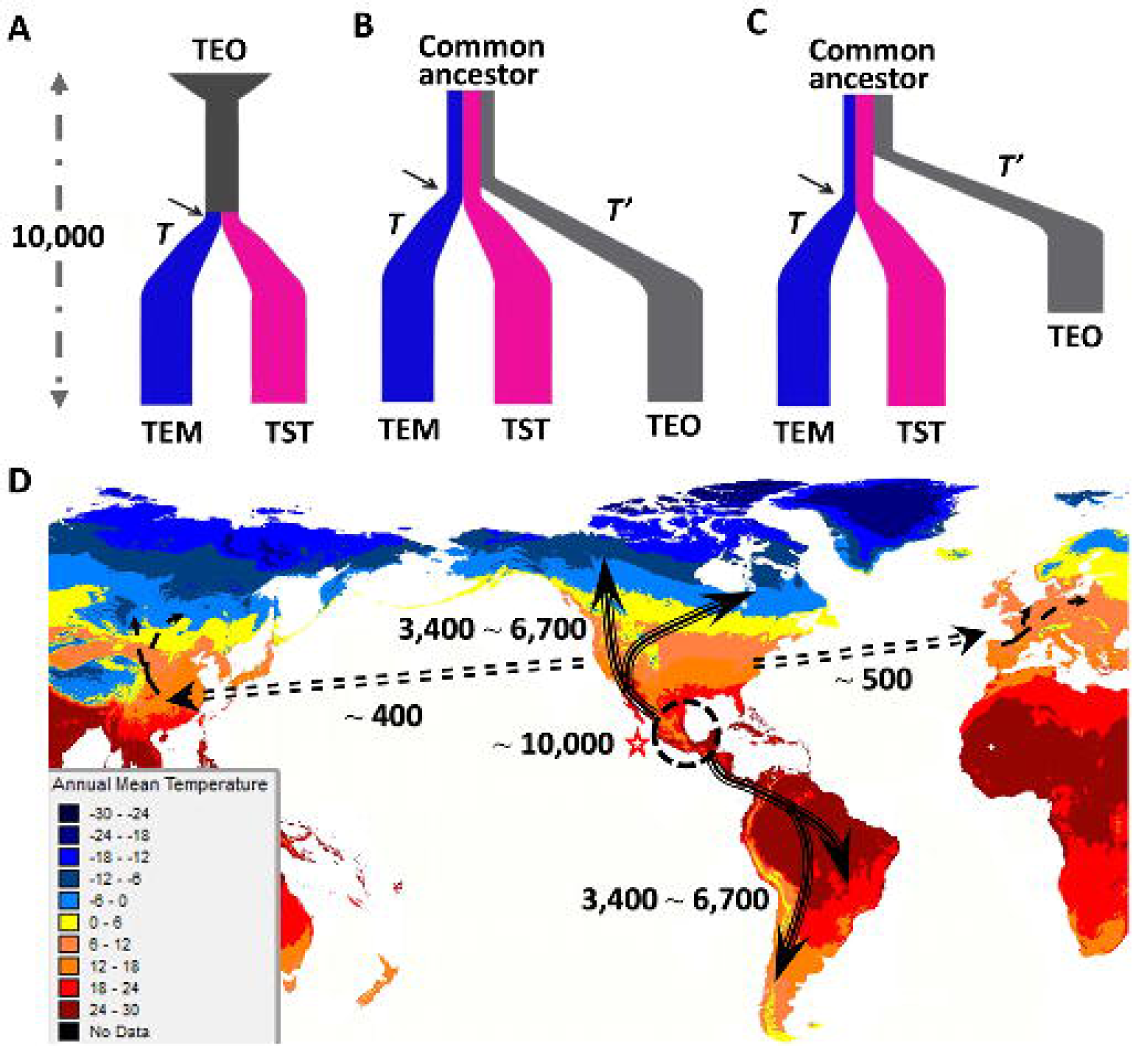
Maize dispersion map and divergence time estimations. (**A**) Divergence time (T) was estimated between temperate (TEM) and tropical/subtropical (TST) maize, using extant teosinte (TEO) lines as the ancestor of maize. (**B**) Divergence time (T) was estimated using a model supposing a common ancestor to TEM, TST, and TEO and equal selection pressure on extant TEO lines (leading to T’ for teosinte) as on TST and TEM. (**C**) Divergence time was estimated between (**A**) and (**B**), with the more likely assumption that TEO lines have experienced a lower level of selection pressure (and smaller T’) than the intensity faced by TEM and TST during the adaptation process. (**D**) Maize dispersion map showing environmental difference in annual mean temperature. Arrows show possible dispersion routes and the numbers beside the routes indicate the likely time (in years before present) when the dispersion happened, and the circle (beside a silver star) is the center of maize domestication (Hufford et al. 2012). Even though there are no inbred lines from South America, the dispersion route to South America was inferred from a previous study (Wallace et al. 2014).

### Genome-wide selection analysis and functional correspondence

The 701 selected regions were compared to 466 and 573 regions identified in domestication and improvement previously, respectively (Hufford et al. 2012; Supplemental Fig. S2). Seven regions were identified in all three processes, and may play a similar role in domestication and post-domestication, contributing to the unique phenotypes of temperate maize. Most adaptation regions did not overlap with domestication and improvement at all, indicating different genomic factors contributing to the phenotype changes during adaptation.

Gene Ontology (GO) annotation of the top 571 candidate genes (see Materials and Methods) within the 701 selected regions reflected genes responding to stress, development, and metabolic processes (Supplemental Table S4). *MFT* (Mother of FT and TFL1, GRMZM2G059358), was identified as a strong candidate gene in adaptation (Fig. 1). It encodes a maize MFT-like protein, which is involved in seed dormancy and germination, a complex adaptation process modulated through a negative feedback loop via ABA (Xi et al. 2010). *MFT* is also known to be involved in control of shoot meristem growth and flowering time (Yoo et al. 2004). Flowering time changes is critical that allow maize to adapt to different environmental conditions such as photoperiod and temperature. Another gene, GRMZM2G360455, an important locus affecting the photoperiod response based on the maize nested association mapping (NAM) population analysis (Buckler et al. 2009), and also containing the “CO/CO-Like/TOC1 conserved site” (CCT), which is always contributed in circadian clock and flowering time (Robson et al. 2001; Griffiths et al. 2003; Cockram et al. 2012), was selected during maize adaptation (Fig. 3A). Metabolic processes influencing nutritionally important traits such as starch content and oil concentration could likely be targets of selection not only during domestication and improvement but also adaptation, as different soil and climate conditions will influence developmental stage of germinating maize. Some genes associated with nutritional traits were selected during the adaptation process in our study (Fig. 1), such as *su2* (GRMZM2G348551), *zpu1* (pullulanase-type starch debranching enzyme1, GRMZM2G158043), and *sdp1* (GRMZM2G087612); these have also been identified as selection targets during domestication and improvement (Zhang et al. 2004; Beatty et al. 1999; Eastmond 2006).

**Figure 3.**
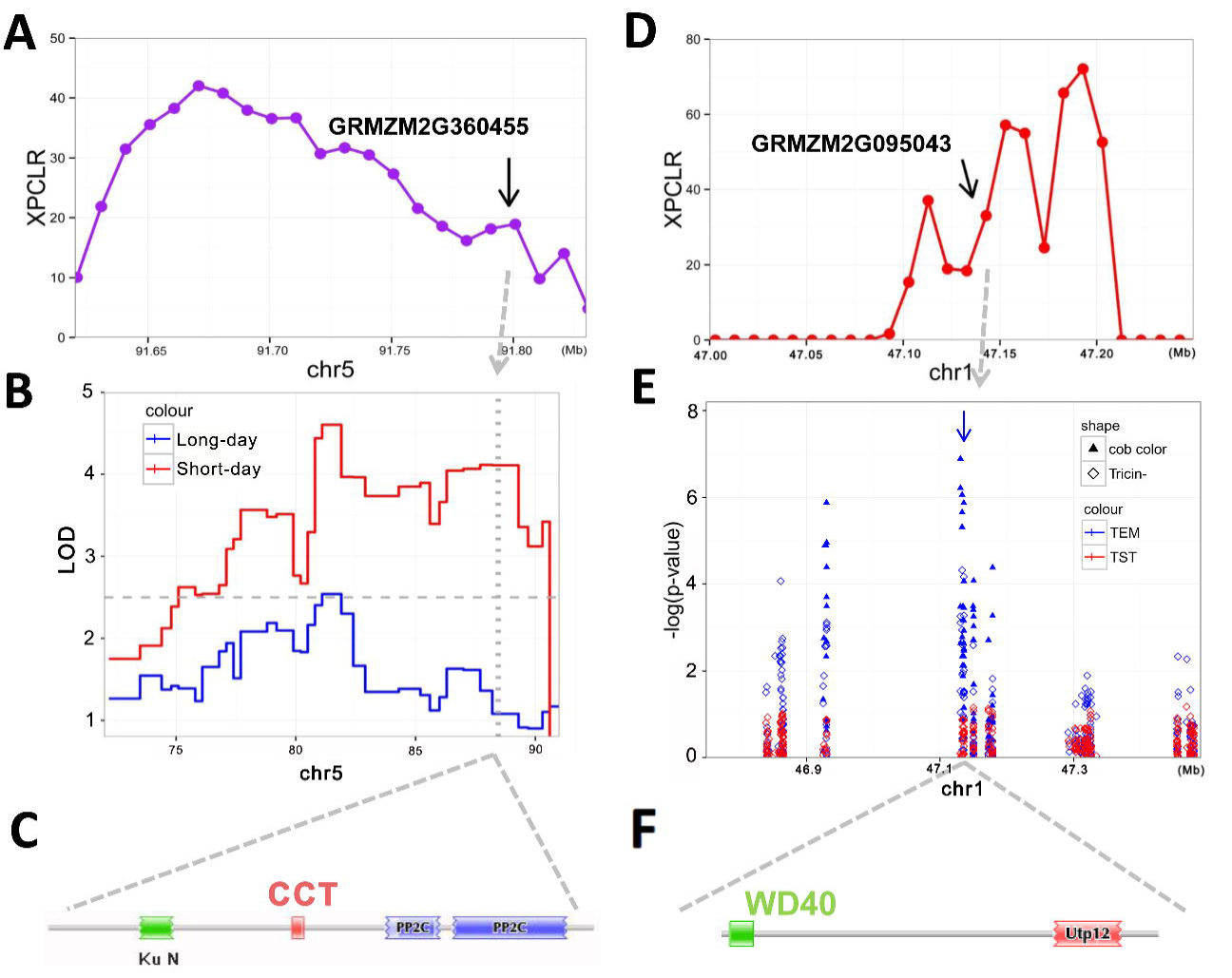
Examples of validation of the function of genes within selected genomic regions, using GWAS, QTL mapping, and gene annotations. (**A**) Gene GRMZM2G360455 was in a selected region on chromosome 5. (**B**) The region which contained GRMZM2G360455 covered a QTL for days to silking (DTS). (**C**) Annotation of GRMZM2G360455 disclosed a CCT domain in this gene which is related to flowering time. (**D**) GRMZM2G095043 was in the selected region on chromosome 1. (**E**) GRMZM2G095043 was strongly associated with cob color and naringenin, which is upstream of the anthocyanin and maysin pathways, and causes changes in cob color. (**F**). Annotation of GRMZM2G095043 indicates that it contains a WD40 domain which could control multiple enzymatic steps in the flavonoid pathway.

Selected metabolic pathways responding to a changing environment are also involved in the adaptation process. Glutathione plays an important role in cellular processes under biotic stress (Dubreuil-Maurizi and Poinssot 2012) and is one such example. Accordingly, *gst35* (Glutathione S-transferase 35, GRMZM2G161891), *gst41* (Glutathione S-transferase 41, GRMZM2G097989) and *gsh1* (gamma-glutamylcysteine synthetase1, GRMZM2G111579), which influence glutathione metabolism, were all within our selected regions (Fig. 1). Traits related to plant architecture and vegetative growth were also selected during adaptation, and their corresponding genes were found in our top XP-CLR hits, including *rs2* (rough sheath2, GRMZM2G403620), essential for normal leaf morphology (Phelps-Durr et al. 2005); *bk2* (brittle stalk2, GRMZM2G109326) affecting mechanical strength of maize by altering the composition and structure of secondary cell wall tissues(Ching et al. 2006), and *apt1* (aberrant pollen transmission1, GRMZM2G448687), affecting the elongation of root cortex cells and pollen tubes during temperature stress (Xu and Dooner 2006) (Fig. 1, Supplemental Table S3).

### Phenomic changes affecting adaptation and validation of selected regions.

Adaptation involves selection of different ecotypes suited to different environments, leading to measurable phenotypic differences between environments. To test this assumption, we collected 662 phenotypic data points including agronomic, yield, seed quality, and seed metabolic traits (Supplemental Tables S5, 6, 7; Wen et al. 2014). The *Q_ST_* – *F_ST_* method (Leinonen et al. 2013), by comparing complex partitioning of variation of quantitative traits and neutral molecular markers (see methods for details), allowed us to distinguish whether the divergent traits are caused by directional selection (*Q_ST_* > *F_ST_*), genetic drift (*Q_ST_* ≈ *F_ST_*), or stabilizing selection (*Q_ST_* < *F_ST_*) (Leinonen et al. 2013). One hundred and thirty traits displayed both divergent patterns suggestive of directional selection across the populations and significant (p≤5.1E-05) differences between the tropical and temperate subgroups (Supplemental Table S8), which were likely to contribute to improved phenotypic performance in temperate conditions.

To test if selected regions contributed to these phenotypic changes, a genome wide association study (GWAS) was performed on the 130 divergent traits with 24,595 SNPs from the selected regions. In total, 345 regions (49.22% of the total) were associated with 100 of the 130 traits (P < 4.06E-05), including three agronomic (days to silking, cob color and kernel color), one amino acid (Ala), and 96 metabolic traits (Supplemental Table S9). The genes identified here undoubtedly represent only a fraction of the total genes selected during adaptation, since many target traits were not measured in this study.

Resistance to new biotic stresses was essential to the adaptation process. Gene GRMZM2G095043 falls within a selected genomic region, and likely contributes to resistance to new pathogens faced by migrating maize. This gene contains a WD40-repeat domain potentially involved in the regulation of the flavonoid pathway (Koes et al. 2005), and showed a strong association with both cob color and Tricin O-pentosyl-O-hexoside (n1570) in temperate lines in this study (Fig. 3D, E, F; Supplemental Table S9). Tricin O-pentosyl-O-hexoside is a flavonoid, and flavonoids are colored compounds that increase insect resistance (McMullen et al. 2009; Falcone et al. 2012).

Quantitative trait locus (QTL) mapping was used to confirm the phenotypic effect of the selected regions. As flowering time changes is representative that allow maize to adapt to temperate environmental conditions such as photoperiod and temperature. Three RIL populations generated by crossing temperate lines with tropical/subtropical lines were used to map flowering time (Supplemental Table S10). The identified QTL were reported in Supplemental Table 7, and many (113, or 62%) overlapped significantly (P < 2.2E-16) with the identified selected regions (Fig. 1). For example, gene GRMZM2G360455, encoding a CCT domain-containing protein was located in the QTL interval for days to silking under short day (tropical) but not under long day (temperate) conditions, and also a candidate response to flowering time of the maize NAM population (Buckler et al. 2009), was identified within a selected region (Fig. 3A, B, C). Further study of this gene may provide a better understanding of maize flowering time and adaptation.

### Significance of transcriptional regulation to adaptation

#### Differential expression

Changes in gene regulation, impacting gene expression level but not gene structure, are fundamental to the evolution of morphological and developmental diversity (Swanson-Wagner et al. 2012; Carroll 2008). The present datasets provide an excellent resource for investigating the contribution of transcriptome regulation to the adaptation process. The coefficient of variation (COV) was similar between tropical and temperate germplasm when considering gene expression of all genes (Fig. 4A), suggesting that most inbreds were probably at the same developmental stage and that no transcriptome-wide changes occurred between tropical and temperate lines. This agrees with a previous study (Swanson-Wagner et al. 2012), indicating that overall changes in expression did not happen during domestication and post-domestication. To study specific transcriptome signal response of individual genes or groups of genes contributing to the adaptation process, *Q_ST_*-*F_ST_* of differentially expressed genes were compared, including single genes differentially expressed under different conditions or in different samples (DE), and genes with altered expression conservation (AEC) (Swanson-Wagner et al. 2012), representing the rewired co-expression of a gene network.

**Figure 4.**
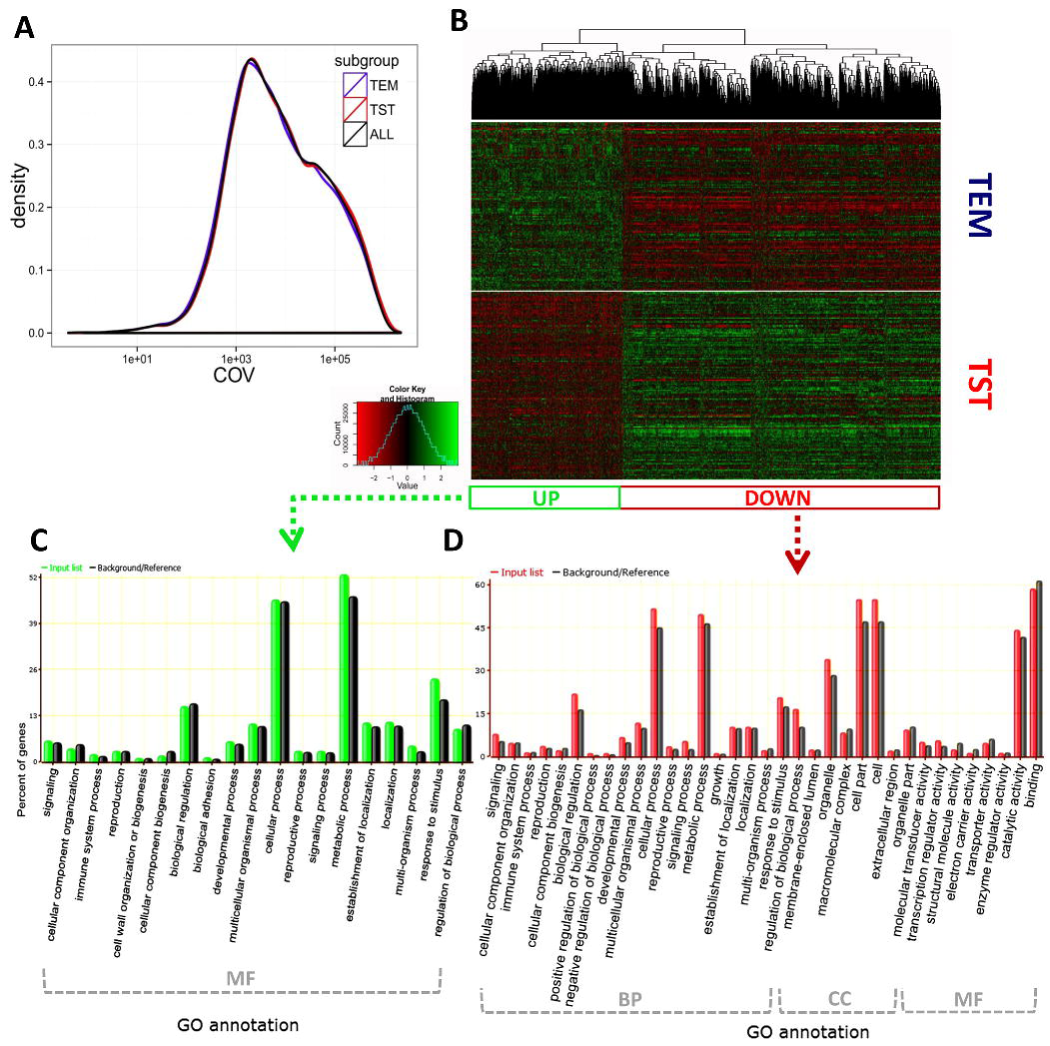
Transcriptional differential expression analysis. (**A**) Density plots for the coefficient of variance (COV) for gene expression levels in all lines (black), temperate lines (blue), and tropical lines (red). (**B**) Genes with significantly different expression were used for hierarchical clustering. (**C**) Enrichment analysis of GO annotation within up-regulated genes in temperate lines relative to tropical lines. (**D**) Enrichment analysis of GO annotation of down-regulated genes in temperate lines relative to tropical lines. Within the specific GO terms in (**C**) and (**D**), MF represents genes of molecular function, CC represents genes with a cellular component, and BP are genes associated with biological processes.

Among 28,769 expressed genes, 2,700 (9.4% of the total) showed significant differential expression (posterior prob. > 0.9999); all exceeded neutral expectation (Supplemental Table S11). Comparing between temperate and tropical lines in the public available database with the same quantitative measurement and DE analysis process, we found a major part of these DE genes were also expressed differently in other tissues, including the shoot apex (Li et al. 2012), (P =1.88E-38) and seeding L3 leaf (Eichten et al. 2013), (P =1.44E-15; Supplemental Fig. S3). This is highly suggestive that most differentially expressed genes identified here were not caused by random environmental and developmental variation and that many of the candidate DE genes continued to be important to adaptive differences later in the development of the mature plant. With more relaxed posterior probability (> 0.999), 14.4% of the total genes showed expression differences, and most were still likely to have been caused by directional selection (Supplemental Fig. S4).

There were 871 up- and 1,829 down-regulated genes in temperate vs. tropical lines (Fig. 4B, Supplemental Table S11). These DE candidates tend to be regulated by distant eQTLs (P=1.74E-4, eQTL data from Fu et al. 2013), and especially for up-regulated (P=9.67E-12) but not for down-regulated (P=0.83) genes in temperate maize. With local (cis-) regulation, expression differences are caused by one regulatory region near each expressed gene, while more distant (trans-) regulation often sees the expression of whole groups of genes regulated by a single genetic factor, causing potential widespread pleiotropic effects. Thus, trans-regulatory mutations seem to be better suited to the changes of complex phenotypes which are governed by the coordinated expression patterns of multiple genes and a single regulator. While some previous studies have emphasized the key role for cis-regulatory evolution (Wray et al. 2007; Stern et al. 2009), our results suggest that trans-regulatory variation could contribute more commonly to adaptive phenotypic divergence.

Gene Ontology (GO) analysis (Fig. 4C, D) of DE genes showed an enrichment of up-regulated genes involving molecular function, especially in catalytic activity, oxidoreductase activity, endopeptidase inhibitor activity and transferase activity (Supplemental Fig. S5). Endopeptidase levels are influenced by stress in plants (Antão and Malcata 2005; Richen et al. 2003) and animals (Karlsson et al. 2006), but clear mechanisms are still unknown. The GO analyses of down-regulated genes in temperate lines revealed enrichment of genes involved in processes such as response to stimuli, metabolism, and regulation (Supplemental Fig. S5). The same GO analyses also uncovered genes involved in cellular components and molecular functions; one down-regulated protein serine/threonine phosphatase involved in both aspects was particularly significant and is associated with biotic and abiotic stress (Antão and Malcata 2005) (Supplemental Fig. S5).

The MADS-box family of transcription factors is important in the evolution of plant architecture and angiosperm inflorescence development, and is frequently identified as targeted regions of selection during the domestication of maize (Zhao et al. 2011). Three MADS-box genes [*zmm5* (GRMZM2G148693), *zmm29* (GRMZM2G152862) and *zap1* (GRMZM2G171365)] were all down-regulated in temperate lines (Supplemental Fig. S6). These genes belong to the MICK^c^ class of the MADS-box gene family and the TM3, GLO and SQUA subfamilies, respectively, and are involved in growth and flower homeotic function (Münster et al. 2002). The circadian clock is vital in flowering time networks which consist of three negative feedback loops. Gene *TOC1b* (GRMZM2G148453) is located in the central loop (Kolmos et al. 2008), and showed down-regulation in temperate lines in the current study (Supplemental Fig. S6). Some domestication and improvement genes (James et al. 1995; Jackson and Hake 1999; Gross and Olsen, 2010) were also identified in the DE analysis, such as *tb1* (teosinte branched 1, AC233950.1_FG002), *su1* (sugary 1, GRMZM2G138060), and *abph1* (aberrant phyllotaxy1, GRMZM2G035688) (Supplemental Fig. S6). The presence of domestication or improvement genes in the adaptation process implies sustaining natural and artificial selection throughout the entire process of maize evolution.

Only 129 genes were identified both in DE and genome selection analysis. Gene Ontology analysis of these genes further indicates a diverse range of biological functions, from response to stimulus to biological regulation and metabolic processes (Supplemental Fig. S7). Since the two data sets have few genes in common, no specific pathways were highlighted in the simultaneous analysis of the two data sets. Additional genes could be found in transcriptome analysis of RNA collected from other tissues and organs, and a more complete annotation of maize genes would increase the number of genes in known pathways, which are currently incomplete.

To seek association between DE genes and the divergent traits, transformed expression values of DE genes were analyzed via DE-GWAS (see Materials and Methods) with the 130 traits that had been found to be significantly different between tropical and temperate lines. Two hundred and forty five DE genes (9.07% of the total) were associated (P<1E-3) with 101 traits, including 75 traits overlapping with those detected by trait-SNP GWAS (Table 1). The 101 DE associated traits included 6 agronomic (cob color, days to silking, days to tasseling, days to pollen shedding, leaf number and kernel color), 5 amino acid (Ala, Arg, Asp, Lys, Gly) and 90 metabolic traits (Supplemental Table S12). Seven of the genes identified as associated in the genomic sequence analysis overlapped with genes identified in the DE-GWAS analysis.

**Table 1.** Summary of SNP-trait and expression-trait association studies

|  | Associated | Agronomic | Amino | Metabolic | Related |
| --- | --- | --- | --- | --- | --- |
|  | traits<sup>a</sup> | traits | acid traits | traits | genes<sup>b</sup> |
| Genome | 100 | 3 | 1 | 96 | 518 |
| Transcriptome | 101 | 6 | 5 | 90 | 245 |
| Overlapped | 75 | 3 | 1 | 71 | 7 |
| Total | 126 | 6 | 5 | 115 | 756 |

<sup>a</sup> represents the divergent traits which were associated with varied markers (SNPs or DE genes) using genome wide association study (GWAS).
<sup>b</sup> represents the closest genes (or within genes) of the significant associated markers.

A series of DE genes (*c2, pr1, a1, bz1, whp1*) were detected that influence cob color via the flavonoid biosynthetic pathway (Sharma et al. 2012) (Fig. 5). Cob color segregated only in the temperate group and is known to be affected by chalcone synthase (CHS) and maysin synthesis, which are thought to be major contributors to corn worm resistance (McMullen et al. 2009; Sharma et al. 2012). As can be seen in Fig. 5, *c2* (colorless2, GRMZM2G422750) is one of the main genes of the CHS pathway and is located upstream of maysin synthesis (Sharma et al. 2012). *pr1* (purple aleurone1 or red aleurone1, GRMZM2G025832) is located downstream of *c2,* encodes a CYP450-dependent flavonoid 3’-hydroxylase required for synthesis anthocyanin, and is involved in naringenin chalcone catabolism (Sharma et al. 2012). *a1* (anthocyaninless1, GRMZM2G026930), located downstream of *c2* and *pr1*, is involved in the production of anthocyanins (Sharma et al. 2012). *c2* and *a1* were significantly associated with flavonoid catabolism (Supplemental Table S12). Kernel color ranked from light to dark (Supplemental Fig. S8A) displayed different distributions between tropical and temperate subgroups (Supplemental Fig. S8B), and was slightly negatively correlated with cob color (Supplemental Fig. S8C, P = 1.31E-05). Kernel color was also affected by DE genes within the CHS pathway and different association patterns were observed between tropical and temperate lines by DE-GWAS analysis (Supplemental Fig. S8D).

**Figure 5.**
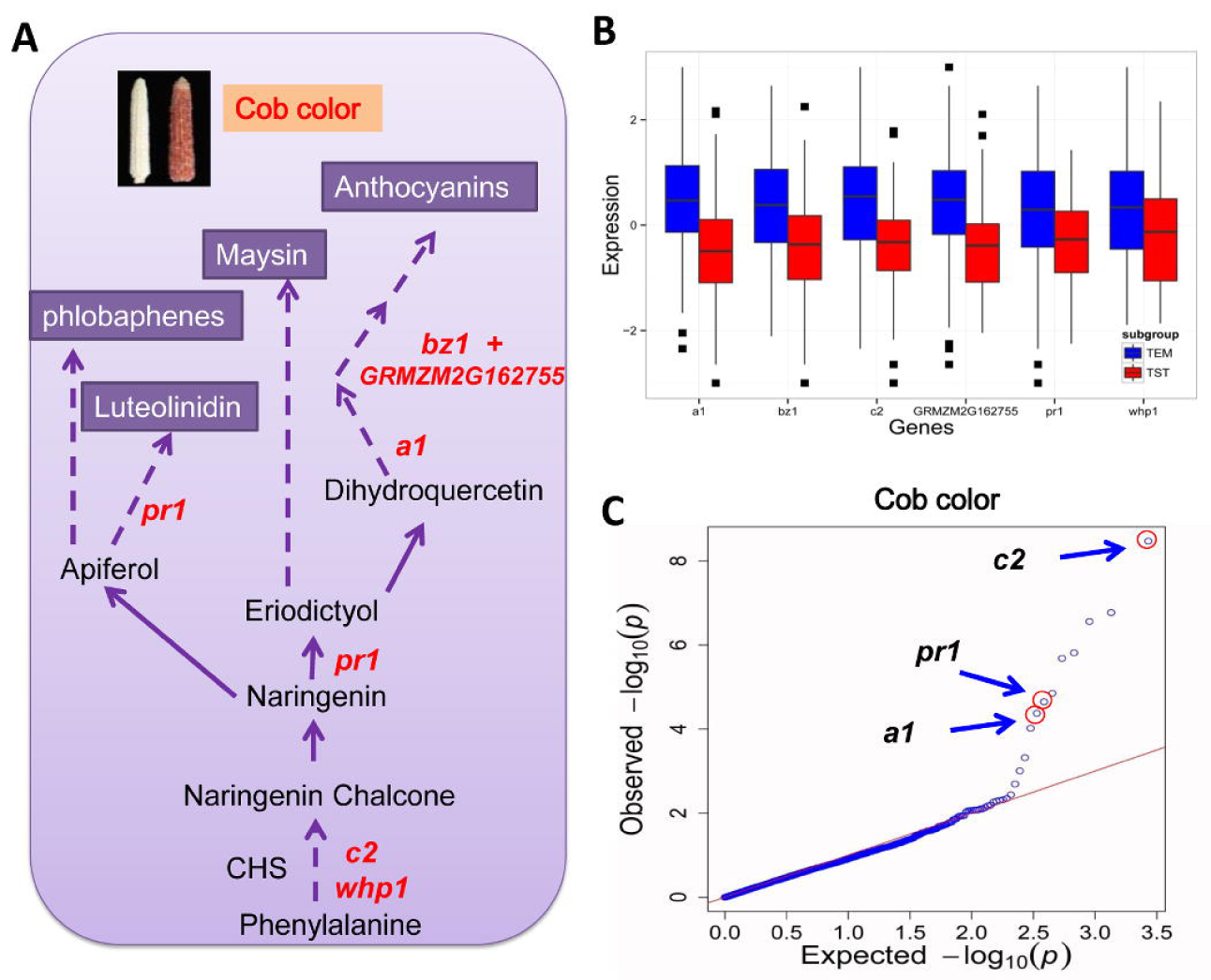
Cob color was influenced by the flavonoid biosynthetic pathway in maize. (**A**) A simplified flavonoid biosynthetic pathway. Genes in red were found to be associated with cob color in this study. (**B**). Genes involved in the flavonoid pathway showed significantly differential expression between temperate and tropical groups. (**C**). Three DE genes (*c2* (GRMZG2G422750)*, pr1* (GRMZG2G025832)*, a1* (GRMZG2G026930)) had significant association with cob color.

### Altered expression conservation

While DE can identify individual differentially expressed genes, altered expression conservation (Swanson-Wagner et al. 2012) (AEC) reflects the relationship of genes with their co-expressed group. In total, 389 genes showed the strongest AEC patterns (expression conservation score >2.5SD; Supplemental Table S13). Further analysis indicated that the average number of genes significantly co-expressed with AEC candidates (as measured by an absolute Pearson correlation coefficient ≥ 0.3) in temperate lines was higher (259 genes) than in tropical lines (147 genes; Supplemental Fig. S9). This indicates that the changes faced in temperate environments may have enhanced interactions between genes in certain pathways. Among the stronger relationships, the number of genes with negative correlations (Pearson correlations ≤ 0) was higher in temperate germplasm as well (24.4% in temperate lines vs. 3.2% in tropical lines). Few genes were co-expressed with the same candidate AEC gene in both temperate and tropical lines (Supplemental Fig. S9), which suggests a rewiring of the regulation networks in the temperate subgroup during adaptation. However, the rewiring appears to have been less dramatic during adaptation since fewer AEC genes were identified for adaptation than for domestication (Swanson-Wagner et al. 2012). In contrast, the number of differentially expressed individual genes found in the adaptation analysis was higher than for domestication (Swanson-Wagner et al. 2012).

As sessile organisms, plants have evolved to integrate endogenous and external information and employ signal transduction processes to allow growth plasticity and survive and thrive in their environments. Essential to this plant-environment interaction are plant hormones, including auxins, ethylene, abscisic acid, and brassinosteroids, which play a key role in plant growth, development, defenses and stress tolerance (Wolters and Jürgens 2009). Plant hormones modulate gene expression by controlling either the abundance of transcriptional factors or repressors, or their activities through post-translational modifications (Dharmasiri et al. 2013). Hormonal cross-talk and interaction with other plant compounds and environmental factors rely on complicated signaling networks. These networks generally intersect in central nodes. Six candidate genes related to plant hormones were observed in our AEC analysis and may act as central nodes in plant hormonal production. Three (GRMZM2G078480, GRMZM5G860241 and GRMZM2G086773) belong to brassinosteroid biosynthesis pathway, one (GRMZM5G864847) was an auxin-responsive Aux/IAA family member, and two (GRMZM2G026131, GRMZM2G390385) were in the pathway producing ethylene biosynthesis from methionine (Supplemental Fig. S10). These candidates, along with their co-expressed genes, showed very different networks in temperate vs. tropical germplasm (Supplemental Fig. S10). This could provide helpful clues to a deeper understanding of the complex relationship between hormones and their contributions to maize adaptation.

Many studies have identified the function of F box receptors in hormone controlling and signaling (Dharmasiri et al. 2005; Kepinski and Leyser 2005; Koops et al. 2011; Shen et al. 2012). AEC analysis also identified an F-box protein (GRMZM2G031958; Jia et al. 2013) displaying different regulation patterns, with a series of 462 and 85 highly co-expressed genes observed in temperate and tropical lines, respectively. Beyond the remarkable difference in network size, several co-expressed enzymes, including cytochrome-c reductases, NADH-ubiquinone oxidoreductases, peroxidases and amidase, and some genes functioned in abscisic acid and IAA biosynthesis and gluconeogenesis were only exists within temperate lines. These results are consistent with the earlier studies that F-box proteins could play a regulatory role in glucose-induced seed germination by targeting ABA synthesis (Song et al. 2012). F-box proteins also play critical roles in seed development, grain filling and response to abiotic stress(Jain et al. 2007) in crop plants, and co-expressed (with our F-box candidate) genes involved in cytokinins degradation, cellulose biosynthesis, and flavonoid and flavonol biosynthesis pathways were observed in the temperate subgroup but not in the tropical. This F-box gene was also highly co-expressed with an ethylene-responsive factor-like protein (GRMZM2G169382), an abscisic stress protein homolog (GRMZM2G044132), a SAUR37-auxin-responsive family member (GRMZM2G045243), another two F-box domain-containing proteins (GRMZM2G116603 and GRMZM2G071705) and several transcriptional factors (zmm29, GRMZM2G152862; ethylene-responsive transcription factors, GRMZM2G474326 and GRMZM2G169382; WRKY1, GRMZM2G018487; WRKY25, GRMZM2G148561; WOX2B, GRMZM2G339751; and more with putative regulation of transcription functions). In addition, the teosinte glume architecture 1 (tga1) protein and an Early Flowering 4 (EF4)-like protein simultaneously showed strong relationships with the target F-box gene. These two proteins have been hypothesized to contribute to maize adaptation to temperate climates in past studies (Khanna et al. 2003; Ducrocq et al. 2008). The identification of the genes encoding these proteins made the network more complex, and further studies are needed. Gene Ontology analysis of all the co-expressed genes in temperate lines revealed an abundance of genes that respond to temperature stimulus and abiotic and other stress (Supplemental Table S14), which were undoubtedly required during maize adaptation to temperate environments.

## Discussion

The process of adaptation was as important a driving factor as domestication in the creation of a geographically diverse crop, allowing maize to spread into a wide range of environments around the world. The impacts of domestication and improvement on the genome and transcriptome have started to be studied (Hufford et al. 2012; Swanson-Wagner et al. 2012), but adaptation has not been as systematically analyzed. In this study, large sets of genomic, transcriptomic and phenomic data were used to analyze the mechanisms of morphological evolution leading to adaptation. Plant response to the environmental change especially stress may have been the key initial step towards adaptation to more extreme latitudes (Fig. 2D). A variety of mechanisms could have contributed to stress response during the maize life cycle, including changes in seed dormancy, germination, plant architecture, flowering time, and optimal utilization of resources (nitrogen, water, etc). In the face of new biotic stresses, a series of resistance mechanisms can also evolve at both the genomic and transcriptomic levels. Traits allowing the plant to successfully respond to stress are precisely those of interest to farmers and plant breeders who have achieved improvement of temperate lines by the selection of such stress tolerance (Tollenaar and Wu 1999).

It has been suggested that studies scanning for positive selection may incur high false-positive rates and can be misleading (Pavlidis et al. 2012). In this study, we use two additional and reliable methods, GWAS and QTL mapping, to provide stronger evidence for the selected regions and a link to the contributory traits. We also identified transcriptomic variants contributing to the adaptation process, including differentially expressed individual genes and evidence for rewiring of co-expression networks. These candidate genes and regions were found to be functionally related to stress response and most were associated with the directionally selected traits. While this study focused on seed transcriptome, the seed-expressed genes and phenotypes provide the first steps towards a systematic study of the adaptation process and inform our understanding of the extent to which transcriptome variation influences the environmental adaptation process.

It has been recently reported that human adaptation is driven primarily by gene expression changes (Fraser 2013). The present study reveals that transcriptome regulation was also prevalent in maize adaptation. Plants live in a dynamic environment, but selection on genomic variation may be too slow to cause the changes that allow the plant to adapt to a rapidly changing environment (Chen 2007). Selection on protein coding sequence variation is risky, as most mutations are harmful or lethal to the organism, and even changes which are beneficial to some of cells or under some conditions may be harmful to other tissues or under other conditions (Ecker et al. 2012). Sufficient protein-coding sequence changes at the genomic level would be unlikely to respond to environmental changes experienced in one or a few generations; however, rapid changes in transcriptome regulation can occur quickly and lead to a rapid phenotypic differentiation (Chen 2007). Transcriptome changes are resource-economical and are frequently associated with temporally and spatially related gene-expression patterns, the effects of which can be limited to specific cells (Carroll 2008; Ecker et al. 2012). Our models, indicating a separation time between temperate and tropical maize of 3,400-6,700 years BP, suggest that a great number of changes took place during a few thousand years. The differences in transcription of individual genes and correlated suites of genes between temperate and tropical maize can be explained by this hypothesis. Some studies (Cortessis et al. 2012; Ptashne 2007) have suggested that epigenetic regulation was the main genetic driver causing regulatory evolution, and that epigenetic modification could also lead to increased mutation rate (Rideout et al. 1990; Schuster-Böckler et al. 2012). Further study and more direct evidence will be needed to better understand the interplay between epigenetic and genetic processes under selection and provide new insights, and possibly new mechanisms, for practical plant improvement.

Although our comprehensive study of genomic, transcriptomic and phenomic variation sheds new light on the process of adaptation in modern maize, much is left to be uncovered. In particular, changes due to adaptation of temperate maize are inferred in the current study by comparing temperate maize with tropical maize grown in northern temperate growing areas. Although similar divergence and changes may have happened as maize migrated to far southern temperate regions, no South American maize was studied here and thus this conclusion remains to be confirmed. In future studies, expression data from more tissues and genotypes need be included and studied under more environmental conditions to allow a finer dissection of genes and mechanisms involved in adaptation. It also should be kept in mind that all the genotypic variation was identified by comparison to the temperate maize reference genome (B73). If a bias was introduced by this method of polymorphism identification, it was not considered in this study. Novel assembly strategies taking into account the variation from more lines could reduce this bias (if present), and exploration of more variant types, such as presence/absence variation (PAV), should also be considered in future studies of the adaptation process. Detailed studies on changes in the transcriptome and in particular, the role of epigenetics, could lead to a clearer understanding of adaptation, possibly leading in turn to more innovative techniques to allow plant breeders to apply native trait variation to maize improvement.

## Methods

### Maize inbred lines and collection of genotypic, phenotypic and gene expression data.

The 368 maize inbred lines included in this study form a global collection including representative temperate and tropical/subtropical inbred lines listed in Supplemental Table S1, and additional information about the lines can be found in previous study (Yang et al. 2011).

The poly(A) transcriptome collected from kernels at 15 days after pollination from all 368 lines was sequenced using 90-bp paired-end Illumina sequencing with libraries of 200-bp insert sizes, and 25.8 billion high-quality reads were obtained after filtering out reads with low sequencing quality. A total of 1.03 million high-quality single nucleotide polymorphisms (SNPs) with a missing data rate less than 60% were used for imputation of missing genotypes. Three pairs of biological replicates (SK, Han21, and Ye478) were used to evaluate the reproducibility of genotyping by RNA-seq. The concordance rates were greater than 99% between each pair of replicates (Fu et al. 2013), indicating that the sequencing and SNP calling procedures were reproducible. In addition, all lines were genotyped via the Illumina MaizeSNP50 array. SNPs generated by RNA-seq also met high concordance rates with the genotypes determined by the MaizeSNP50 BeadChip (Fu et al. 2013). Both RNA-seq and MaizeSNP50 data sets were merged to obtain a total of more than 1.06 million SNPs; a final data set of 0.56 million SNPs with a MAF of greater than 5% was produced and more detailed information can be found in a previous study (Fu et al. 2013).

To quantify the expression of known genes with annotation in the B73 reference genome (filtered-gene set, version 2, release 5b), a total of 28,769 genes corresponding to mapped sequence reads in more than 50% of the inbred lines were compiled, and each gene averaged more than 1.5K reads. Read counts for each gene were calculated and scaled according to RPKM (reads per kilo base of exon model per million mapped reads). After RPKM normalization, all genes with a median expression level greater than zero for each sample were included, and the overall distribution of expression levels for each gene was normalized using a normal quantile transformation (Fu et al. 2013). The same three pairs of biological replicates were shown to share most compiled genes (average 95.71%) with high concordance (average person’s r=0.87, (Supplemental Fig. S12) in expression quantification between each pair of replicates. More details on library construction, sequencing, SNP detection, genotype imputation, positive control of SNP accuracy and quantile normalization of expression is described in Fu et al. (2013).

To obtain agronomic traits (reported in Supplemental Table S5), all inbred lines were planted with randomized complete experimental design by single replication in 2010 in four locations (Honghe autonomous prefecture, Yunan province; Sanya, Hainan province; Wuhan, Hubei province; Ya’an, Sichuan province) and 2011 in three locations (Chongqing; Hebi, Henan province; Nanning, Guangxi province). The seven different locations ranged from 18 to 35 degrees north in latitude and from 102 to 114 degrees east in longitude. Kernel color was measured in five additional trials over two years (Sanya, Hainan province in 2011; Honghe autonomous prefecture, Yunan province in 2012; Chongqing in 2012; Wuhan, Hubei province in 2012; Hebi, Henan province in 2012). All agronomic traits were measured and the Best Linear Unbiased Predictor (BLUP) values from different environments and years were used for final analysis (Supplemental Table 2). Maize kernels from each entry in the panel planted in Chongqing in 2011 were collected to quantify amino acid content (Supplemental Table S6), and samples from the panel planted in Yunnan (2011) and Hainan (2010) were harvested to measure metabolic traits. Only the metabolites measured in at least two of the three experiments and showing high correlation (P value<0.05) in the two experiments were retained to calculate BLUP values for the next analysis (Supplemental Table S7).

### Population structure of 368 inbred lines

The STRUCTURE software package (Pritchard et al. 2000) was used to analyze the population structure and the EIGENSOFT analysis package (Patterson et al. 2006; Price et al. 2006) was used to run a Principal Component Analysis (PCA) of the panel used in the present study (Supplemental Fig. S13). Considering the computation time, a SNP marker subset was used for inferring population structure; the subset of 14,685 SNPs was created by removing adjacent SNPs within 50Kbp intervals. All lines were divided into three subpopulations corresponding to stiff stalk (SS), non-stiff stalk (NSS) and tropical and sub-tropical (TST) clusters by STRUCTURE, using a probability of inclusion into each cluster of greater than 0.65(Yan et al. 2009). In the final analysis, there were 133 temperate (103 NSS + 30 SS) lines, 149 tropical (TST) lines, and 86 mixed lines (Supplemental Table S1, Supplemental Fig. S13). The mixed were excluded from further analyses to allow a clearer comparison between tropical and temperate features. The population structure was similar to the analysis of the same lines reported previously using 1536 SNPs (Yang et al. 2011).

### Measuring the genomic changes occurring during the maize adaptation process

To evaluate changes in the maize genome due to adaption, population genetics statistics including π (Tajima 1983) and *F_st_* (Nei 1977) were calculated within the differently adapted maize groups. *F_st_* was estimated as follows:

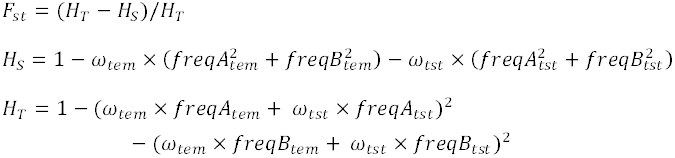

Where *H_S_* refers to heterozygosity within subpopulations, and *H_T_* refers heterozygosity in the overall population. The variable *ω* refers to proportion of subpopulation based on size, and *freqA/freqB* is the frequency of allele A or allele B in each subpopulation. The Synbreed package (Wimmer et al. 2012) of the R-project statistical package (http://www.R-project.org) was used to compute the linkage disequilibrium (LD) coefficient, r^2^. The ggplot2 package of R (Wickham 2009) was used to plot LD decay, as well as all visualization in this study (except as noted, below).

To identify regions of the maize genome that have undergone selection during the process of adaption from tropical to temperate climates, a cross-population composite likelihood approach (Chen et al. 2010) (XP-CLR) was used. The following parameters were applied to implement the XP-CLR test: the size of the window was 0.005 Morgan; the maximum number of controlled SNPs within a window was 100; the spacing between two grid points was 1000 bp; and a corrLevel value of 0.7 was used as a down-weighted criterion in the weighted composite likelihood ratio test. Adjacent 10kb-windows from the top 20% of the XP-CLR results were merged into larger regions, according to Hufford et al. (2012), and only one window lower than 20% was retained for each region. Regions in the highest 10% of the mean region-wise XP-CLR values were regarded as having undergone selection. Gene sequences closest to the maximum XP-CLR value were designated as the most likely candidate genes for selection, while others within each selected region were considered as possible selected genes. The linkage map used in the XP-CLR analysis was constructed using an RIL population generated from the cross of B73 × BY804 with 197 individuals described previously (Chander et al. 2008) and the maize SNP50 chip (Ganal et al. 2011) was used to re-genotype the RIL population with 15,285 polymorphic SNPs. The distance between unmapped SNPs was estimated based on the constructed linkage map and B73 reference genome (version 2).

### Estimating the relative divergence time between temperate and tropical lines

Assuming a time scale of teosinte/maize divergence of about 10,000 years, an *F_ST_*-based approach (Schlebusch et al. 2012) was used to estimate a relative divergence time between temperate and tropical subpopulations, under the assumptions of no genetic drift, no change in effective population size, and equal generation times in each lineage. Although violations of these assumptions are probable and may reduce the accuracy of the estimated divergence times, the estimated divergence times will still be useful for the purposes of this study and in the general study of maize evolution. We combined our SNP data with the maize HapMapII (Chia et al. 2012) data and retained all the SNPs from teosinte (Chia et al. 2012) and maize with the same loci, consistent alleles, and missing ratio of alleles less than 20% to calculate *F_st_* between temperate maize (TEM) and teosinte (TEO, 0.0668), and between TEM and tropical/subtropical maize (TST, 0.0453). Divergence time (T, measured in units of 2*Ne* generations where *Ne* is the e ective number of diploid individuals) was calculated using *F_st_* as according to the following formula (Schlebusch et al. 2012):

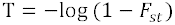

Most of the SNP genotypes were located within expressed sequences, but the results should not be affected by this, assuming similar biases between the two comparisons and sufficient numbers of markers to smooth out unequal biases due to potential unequal selection pressures on some loci. Different models were proposed to improve estimation of the relative divergence time. Assuming the teosinte lines from which we extracted SNPs were indeed the primitive ancestors (or contained the same sequence diversity within them as the actual ancestors), then *T_TEM-TEO_* = 10,000 *years*; (Fig. 2A) and we can calculate the divergence time between TEM and TST using the following formula:

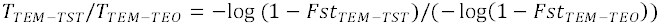

This resulted in a divergence time of *T_TEM-TST_* =6,700 years BP. Assuming further that the teosinte lines have undergone a similar selection pressure (Fig. 2B), the analogical formula can be used to calculate *T_TEM-TST_* divergence time as 3,400 years BP. However, because it is more likely that the teosinte lines have experienced a selection level that is not as strong as that of the adaptation process (Fig. 2C), the true divergence time of adaptation would fall between the two estimated times. Bioclimatic variables data from WORLDCLIM database (Hijmans et al. 2005) and the DIVA-GIS (Hijmans et al. 2012) software was used to map the stress (annual mean temperature) faced by maize during the adaptation process (Fig. 2D).

### Analysis of directional selection of phenomic divergence

*Q_ST_* *- F_ST_* comparison provide us with a method to distinguish population differentiation of complex polygenic traits as having been caused by natural selection or by genetic drift (Leinonen et al. 2013). *Q_ST_* was estimated for all phenomic traits including expression traits as follows:

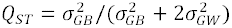

Where 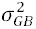 refers to the genetic component of variance among subpopulations, and 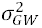 is the average component of variance within each subpopulation.

To accurately estimate the distribution of mean *F_ST_* among tropical and temperate subpopulations, 10,000 SNPs were chosen randomly from the entire SNP data set and the calculation was repeated 1,000 times (Supplemental Fig. S14). By applying a strict outlier definition, we employed a 99% confidence interval (0.025∼0.0273) for the *F_ST_* distribution, ensuring a more correct comparison of *Q_ST_*-*F_ST_*. A *Q_ST_*>*F_ST_* value was regarded as proof of trait divergence outstripping the expectation of a neutral state and thus of a strong directional selection signal (Leinonen et al. 2013).

### Differential expression analysis

Cyber-T (Baldi and Long 2001), a regularized t-test method that also contains statistical inferences on experiment-wide false positive and negative levels based on the modeling of p-value distributions, was used on normalized expression data (Fu et al. 2013) for the identification of statistically significant differentially expressed genes. Posterior Probability of Differential Expression (PPDE) *≥*0.9999 was determined to identify differentially expressed (DE) genes in our study.

### Characterization of genes displaying altered expression conservation

To identify which genes show Altered Expression Conservation (AEC), a statistic which reflects the co-expression of genes in a gene network (Swanson-Wagner et al. 2012), the expression data was divided into 2 matrices (*E^tst^* and *E^tem^*) based on adaptation (tropical and temperate) and a co-expression network was created for each matrix 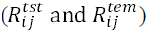. The hmisc package in R (Harrell 2012) was used to calculate the Pearson correlation coefficient between each pair of gene expression values within each subset, (TEM or TST). Hmisc is an efficient algorithm for calculations in very large data sets, and is calculated as:

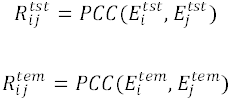

for i, j = 1, …, 28,769. Thus, 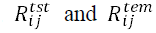 are square matrices with the same dimensions (28,769 × 28,769). Each value in the two matrices represents an edge weight in the co-expression network and is the measured similarity between expression profiles of paired genes. Although the two matrices have identical dimensions, the distribution of values in each matrix may differ because of unequal sample sizes. Thus, to compare the two co-expression networks more accurately, the distributions were normalized by subtracting the mean and dividing by the SD to obtain a standard normal distribution. An expression conservation (EC) score was calculated as the Pearson correlation coefficient between gene profiles in the two co-expression networks as described by Swanson-Wagner *et al*. (2012):

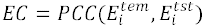

Where 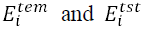 were represented by the i-th rows in the temperate and tropical co-expression matrices, respectively. AEC genes were selected using a *z* score (Swanson-Wagner et al. 2012) to calculate each EC value as follows:

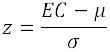

Where μ and σ were the mean and SD of all the gene’s EC scores. The *z* score cutoff of altered expression conservation values of genes was set at ≤ −2.5. The software Circos (Krzywinski et al. 2009) was used for visualizing the results in a circular layout.

### Candidate gene annotation and GO enrichment analysis

To more fully explore candidate genes’ functions, the annotation resources of maizeGDB (Lawrence et al. 2008) and the InterPro (Zdobnov and Apweiler 2001) database were integrated into the analyses. Gene Onotology (GO) enrichment analysis was maintained by AgriGO (Du et al. 2010) with the Fisher statistical test method (P <=0.05) and Yekutieli muti-test adjustment method (FDR<=0.05).The GOSlimViewer of AgBase (McCarthy 2006) was used to implement GO slim analysis, and the updated GO items of the maize genome were downloaded from Ensembl BioMart (Kinsella et al. 2011) on April 4th, 2013.

### Association and QTL mapping analysis

SNPs within regions that have experienced selection according to the XP-CLR approach (24,595 in total) were used in an association analysis with the 111 different traits using the software package GAPIT (Lipka et al. 2012) and a compressed mixed linear model. The cutoff for significance for associations was set at 1/n (n is the number of SNPs used, p <4.06E-05). To better compare trait-SNP GWAS results with the genes identified following DE analysis, the DE identified genes were transformed prior to analysis into a discontinuous pseudo genotype with two alleles, one if expression of the gene for a given line was higher than the median of all lines, and the other if it was lower. Three recombinant inbred line (RIL) populations were used to map QTL: BY population: BK (an inbred selected from a tropical landrace with very big kernel size) × Yu87-1 (an elite inbred with tropical background used frequently in Chinese breeding programs), SZ population: SK (an inbred selected from a tropical landrace with very small kernel size) × Zheng58 (an elite inbred used frequently in Chinese breeding programs), and YZ population: Yu87-1 (an elite inbred with tropical background used frequently in Chinese breeding programs) × Zong3 (an elite inbred used frequently in Chinese breeding programs). The QTL were mapped for three flowering times traits: days to tasseling (DTT), measured as the number of days from planting to 50% male flower appearance; days to silking (DTS) or the number of days from planting to 50% female flower appearance; and days to anthesis (DTA), the number of days from planting to 50% male flower pollen shed. Mapping was done using WinQTLcart (Wang et al. 2011) and a LOD threshold value for significance set to 2.5. The flowering time traits are an indication of adaptation and thus used as a proxy for that trait.

### Comparison with DE gene sets from other tissues

Li et al. (Li et al. 2013) conducted RNA-seq on the shoot apex from 2-wk-old seedlings of the NAM founders. We downloaded the raw data and quantified with the same procedure used in the present study (Fu et al. 2013) and average expression level of the two runs of each line was used. The third leaf (L3) after germination of the NAM founders was also RNA-sequenced by the Springer laboratory, and the RPKM results were obtained by the author (Eichten et al. 2013). These two studies provided different tissues and different genetic backgrounds compared to the current study; these were used to test if genes identified with differential expression in the current study are tissue-specific or if, as assumed in the current study, will prove to have lasting effects during maize development and maturity. Furthermore, consistency of expression differences in different populations can be validated. Genes with zero mapped reads in more than 60% of the inbred lines were excluded and the overall distribution of expression levels for each remaining gene was normalized using the normal quantile transformation (Fu et al. 2013). Lines with unmixed temperate (NSS+SS) and tropical backgrounds were used to call DE genes (P value < 0.05) with the same method (Fu et al. 2013). To test if the number of overlapping DE genes (Supplemental Fig. S3) is significant between the current study and the two published experiments, random subsamples of genes were chosen from each comparison pair (using same the number for each subset as the number of DE genes identified by each study); this simulation was repeated 10,000 times to create a distribution of random overlap DE comparisons (Supplemental Fig. S3). This distribution was used to test if the observed number of DE overlaps is in accordance with the simulated normal distribution.

## Data access

There is no newly generated data in this study. All the data used in this study were published previously, and can be found in the Methods section.

## Acknowledgements

We appreciate the helpful comments on the manuscript from Dr. Major M. Goodman, and Dr. Nathan M. Springer for providing normalized RPKM expression dataset. We thank the students and colleagues from J. Yan’s present laboratory as well as his former laboratory at the China Agricultural University; and the X. Yang and J. Li laboratory at the China Agricultural University for collection of phenotypic data. We thank Dr. C. Bedoya for references on archeological studies in Mexico. This research was supported by the National Natural Science Foundation of China (31123009, 31222041), the National Basic Research and Development Program of China (2011CB100105), the National Hi-Tech Research and Development Program of China (2012AA10A307) and the Fundamental Research Funds for the Central Universities (Program No.2013SC01).

## Author contributions

J.Y. designed and supervised this study. H.L., X.W., W.W., J.L., M.D., H.T., and Q.P. performed the experiments. H.L., X.W. and M.J. performed the data analysis. X.Y. and X. D. contributed new protocols. H.L., X.W., M.L.W. and J.Y. prepared the manuscript, and all the authors critically read and approved the manuscript.

